# Flying In-formation: A computational method for the classification of host seeking mosquito flight patterns using path segmentation and unsupervised machine learning

**DOI:** 10.1101/2021.11.24.469809

**Authors:** Mark T Fowler, Anthony J Abbott, Gregory PD Murray, Philip J McCall

**Author notes:** Address for correspondence: Mathematical modelling, Vector biology.

## Abstract

The rational design of effective vector control tools requires detailed knowledge of vector behaviour. Yet, behavioural observations, interpretations, evaluations and definitions by even the most experienced researcher are constrained by subjectivity and perceptual limits. Seeking an objective alternative to ‘expertise’, we developed and tested an unsupervised method for the automatic identification of video-tracked mosquito flight behaviour. This method unites path-segmentation and unsupervised machine learning in an innovative workflow and is implemented using a combination of R and python. The workflow (1) records movement trajectories; (2) applies path-segmentation; (3) clusters path segments using unsupervised learning; and (4) interprets results. Analysis of the flight patterns of *An. gambiae* s.s., responding to human-baited insecticide-treated bednets (ITNs), by the new method identified four distinct behaviour modes: with ‘swooping’ and ‘approaching’ modes predominant at ITNs; increased ‘walking’ behaviours at untreated nets; similar rates of ‘reacting’ at both nets; and higher overall activity at treated nets. The method’s validity was tested by comparing these findings with those from a similar setting using an expertise-based method. The level of correspondence found between the studies validated the accuracy of the new method. While researcher-defined behaviours are inherently subjective, and prone to corollary shortcomings, the new approach’s mathematical method is objective, automatic, repeatable and a validated alternative for analysing complex vector behaviour. This method provides a novel and adaptable analytical tool and is freely available to vector biologists, ethologists and behavioural ecologists.

**Author summary:** Vector control targets the insects and arachnids that transmit 1 in every 6 communicable diseases worldwide. Since the effectiveness of many vector control tools depends on exploiting or changing vector behaviour, a firm understanding of this behaviour is required to maximise the impact of existing tools and design new interventions. However, current methods for identifying such behaviours are based primarily on expert knowledge, which can be inefficient, difficult to scale and limited by perceptual abilities. To overcome this, we present, detail and validate a new method for categorising vector behaviour. This method combines existing path segmentation and unsupervised machine learning algorithms to identify changes in vector movement trajectories and classify behaviours. The accuracy of the new method is demonstrated by replicating existing, expert-derived, findings covering the behaviour of host-seeking mosquitos around insecticide treated bednets, compared to nets without insecticide. As the method found the same changes in mosquito activity as previous research, it is said to be validated. The new method is significant, as it improves the analytical capabilities of biologists working to reduce the burden of vector-borne diseases, such as malaria, through an understanding of behaviour.

## Introduction

Vector-borne diseases (VBDs) are illnesses caused by Protozoa, viruses and nematodes and transmitted by infected arthropods, such as mosquitoes and ticks. VBDs threaten 80% of the planet’s population, and are responsible for an estimated 17% of all human communicable diseases and over 700 000 deaths annually [1–3]. Many effective strategies to reduce the burden of VBDs target the arthropod vector. Such an approach involves the development and use of interventions that control or exploit vector behaviour and prevent human contact with pathogens. For example, tools that exploit a vector’s host-seeking behaviour include decoys or targets for *Glossina sp*. (tsetse fly, vectors of human animal trypanosomiasis) [4,5] and insecticide-treated bednets (ITNs) for *Anopheles sp*. (the mosquitoes that transmit malaria) and *Aedes sp*. (the principal vector for dengue fever) [6]. Significantly, although these devices are now essential tools for their respective disease control or elimination programmes in sub-Saharan Africa [7], both continue to undergo further research to improve their performance and applicability [8,9]. For example, efforts to improve ITNs have entailed analysis of mosquito net responses through the segregation of flight paths around a human-baited bednet into distinct movements and behaviours. These behaviours were based on flight characteristics detected and defined by the researchers and interpreted as responses to the human, the net itself and/or the presence of any insecticide treatment on the net [9–11]. However, investigations into distinguishing, defining and classifying vector behaviour in such contexts are still principally based on researchers’ expertise and experience with the target species’ biology and ethology [9,12–14]. Nevertheless, reliance on such a solely subjective method is problematic. Expert knowledge is intrinsically inefficient to apply at scale, it is domain-specific, subject to cognitive biases and constrained by the physical limits of human perception [15–17].

Much of this subjectivity can be eliminated by the application of objective computational processes. These processes have the potential to analyse the movement paths of arthropod vectors to isolate and define behaviours, but do so in an automated, repeatable and objective way. Computational algorithms that can investigate animal movement and behaviour in this manner are already available. More specifically, Behavioural Change Point Analysis (BCPA) is a form of path segmentation that splits movements into distinct behavioural ‘bursts’ at significant changes in activity [18–20], thereby isolating movements. For example, BCPA has been used to identify the timing of animal movements [21,22] and to quantify animal behavioural shifts from seasonal environmental changes [23,24]. Secondly, unsupervised machine learning is a statistical approach that can identify hidden patterns present within datasets and has the potential to cluster, and therefore define, movements. These clustering algorithms group datapoints into distinct collections based on any latent structure present within data [25,26] and have been used to identify patterns in neuronal ensembles in the brain [27] and to identify subgroups within patient populations [28]. However, BCPA and unsupervised machine learning are yet to successfully classify insect behaviour from movement trajectories alone. Their application has been restricted in this context, as path segmentation can only identify changes in behaviour rather than behaviours themselves [18], while clustering requires a sufficiently high signal-to-noise ratio to be successful (something raw movement trajectories do not possess) [26].

This study proposes and tests that a solution to the problem of a total subjective base for the classification of vector behaviour is possible by combining the above two identified computational processes. That is, path segmentation and unsupervised machine learning can be brought together to discriminate and categorise vector movements into distinct behavioural modes. However, the combination and application of these algorithms requires a workflow to collect, prepare and analyse trajectories. In this report, we present, describe and test such a workflow. This is a novel method that was devised to support complex behavioural analyses, specifically concerning resource location by mosquitoes, in which: (1) detailed spatial and time-series data covering the movement trajectory of a vector in a domain-specific setting is collected [10,29,30] (Fig 1A); (2) movement trajectories are segmented into behavioural ‘bursts’ through BCPA [19,20] (Fig 1B); (3) these behavioural ‘bursts’ are grouped through an optimised clustering algorithm [31,32] (Fig 1C); and (4) results are interpreted through the analysis of descriptive statistics and examination of representative samples [33–35] (Fig 1D).

**Fig 1.**
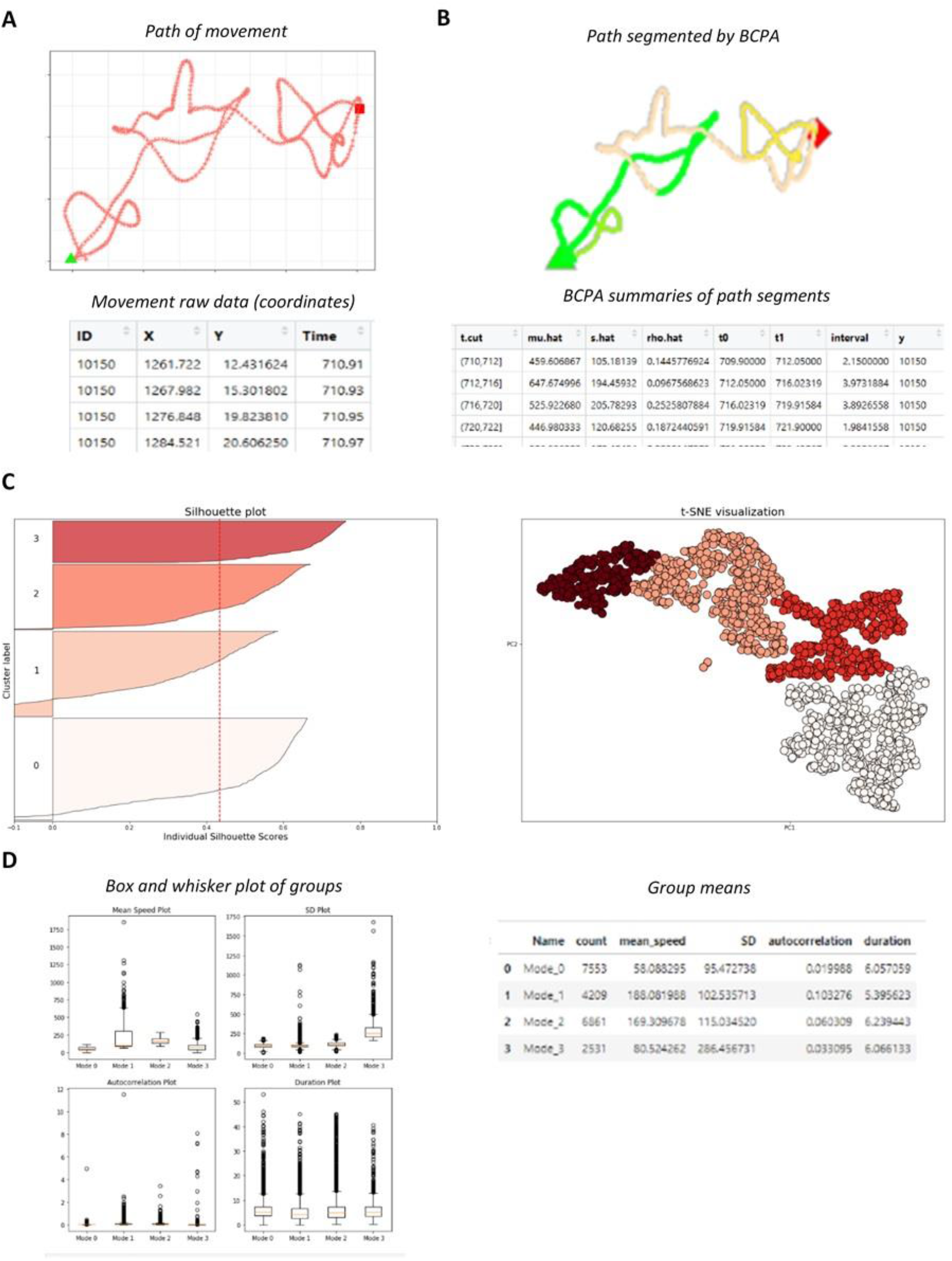
Example of workflow process. (A) Time series data detailing a single vector movement trajectory. For each observation that comprise the movement an identifier, an x-coordinate, a y-coordinate and a time are required. The triangle is the start of the movement, the square the end of the movement. (B) BCPA is used to segment the movement into distinct behaviours, here based on significant changes in persistence velocity. Three significant changes in persistence velocity are identified in this example, giving four tokens of behaviour. BCPA segmentation produces a data frame summarising each phase. (C) The movement segments are grouped using the optimum clustering algorithm and initial parameters, as defined by internal validation. Clustering is internally validated through silhouette score, silhouette plot and manual inspection of a t-SNE visualisation. (D) A label is attached to behavioural groups by interpreting the clustering results. Interpretation is systemised through analysis of group statistics and examination of representative examples from each cluster (i.e., those found at the centre of each groups’ t-SNE plot).

Combining path segmentation and unsupervised machine learning into a single unique workflow provides a novel method to overcome the inherent subjectivity and perceptual limitations of any investigator-led alternative. In a first application of the method, we analysed the flight paths of the primary African malaria vector mosquito, *An. gambiae* s.s., during host location around an ITN with a single human occupant, recorded under experimental conditions in the laboratory. We report that the new workflow distinguished four behavioural types that varied in frequency depending on net treatment. These findings corresponded well with those in a previous investigator-led interpretation [9], but were achieved in a more objective, repeatable manner.

## Results

To assess the accuracy of the new workflow, we applied the method to the activity of *An. gambiae*, a principal vector of malaria in sub-Saharan Africa, around either an insecticide-treated net (as approved by the World Health Organisation, hereafter ‘treated’) or an untreated polyester net (‘untreated’). A strain of mosquito susceptible to all insecticides, Kisumu, was used in both the untreated and treated arms of the experiment. The results of this application were then compared with those from a previous, expert-derived, study to validate the accuracy of the workflow.

### Data acquisition, cleaning and assessment

Activity rates, based on observations from the raw data, were found to be much higher around an untreated net, with the number and length of movements significantly lower when an ITN was used (Table 1). When the autocorrelation of the datasets was assessed, it was found that movement velocity was positively autocorrelated through 50 time-lags in both the untreated (Fig 2A) and treated (Fig 2B) data. Accordingly, the data were taken to be suitable for analysis.

**Fig 2.**
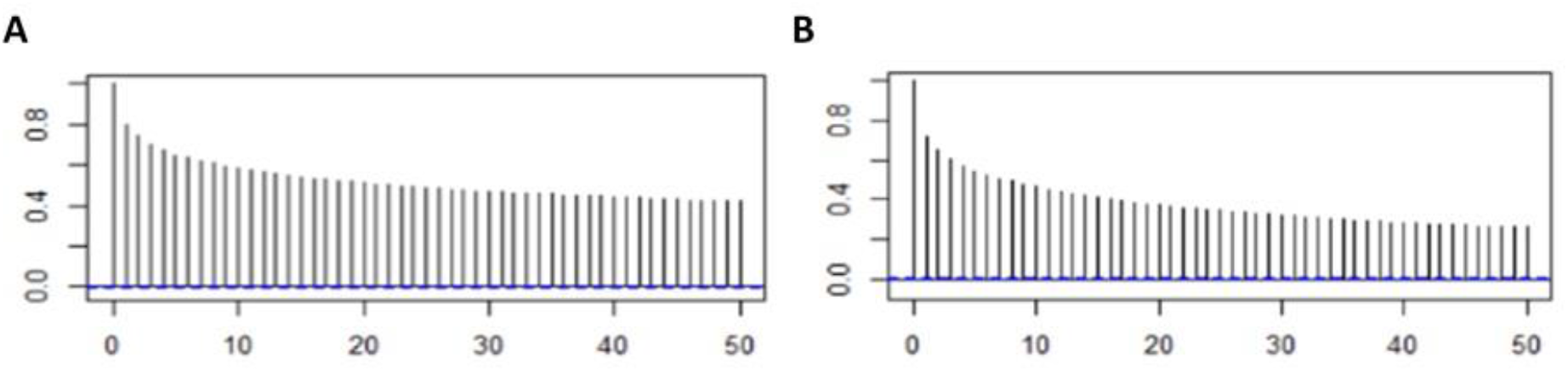
Correlograms. (A) Untreated velocity correlogram. Movement speed is autocorrelated with its recent past (through a maximum of 50 lags). This association becomes weaker as the lag increases (from 0.8 at lag 1 to 0.5 at lag 50). (B) Treated velocity correlogram. Movement speed is autocorrelated with its recent past (through a maximum of 50 lags). This association becomes weaker as the lag increases (from 0.7 at lag 1 to 0.3 at lag 50).

**Table 1.** Untreated and Treated tracking and segmentation figures for evaluation. All BCPA set to detect a significant change in persistence velocity (‘Velocity*cos(Turning Angle)’), using a window size of 30, a window step of 1, a sensitivity value of 2 and a cluster width of 1.

|  |  | Untreated | Treated |
| --- | --- | --- | --- |
| Tracking - Raw | Replicates | 5 | 5 |
|  | Total Length | 10 hrs | 10 hrs |
|  | Strain | Kisumu | Kisumu |
|  | Observations | 3 514 999 | 491 708 |
|  | Trajectories | 9 076 | 1 472 |
|  | Max trajectory length | 215.3 secs | 124.0 secs |
|  | Mean trajectory length | 7.4 secs | 6.7 secs |
|  | Mode trajectory length | 0.5 secs | 0.5 secs |
| Tracking - Cleaned | Observations | 3 320 055 | 453 201 |
|  | Trajectories | 5 475 | 798 |
|  | Max trajectory length | 215.3 secs | 124.0 secs |
|  | Mean trajectory length | 12.1 secs | 11.4 secs |
|  | Mode trajectory length | 2.2 secs | 2.1 secs |
| Segmentation (BCPA) | Total phases | 33 350 | 3 979 |
|  | Max trajectory phases | 61 | 40 |
|  | Mean trajectory phases | 6 | 5 |
|  | Mode trajectory phases | 2 | 2 |

### Path segmentation

Results of the path segmentation are found in Table 1. Activity of *An. gambiae* s.s. was again found to be significantly higher in the untreated trial.

### Clustering

Internal validation indicated that the optimal algorithm and parameters to cluster both the untreated and treated data was an agglomerative clustering algorithm using Ward’s method for linkage and four clusters.The untreated clustering produced a silhouette score of 0.36 (Fig 3A), while the treated grouping’s silhouette score was 0.41 (Fig 3B). Results of this analysis are shown in Table 2.

**Fig 3.**
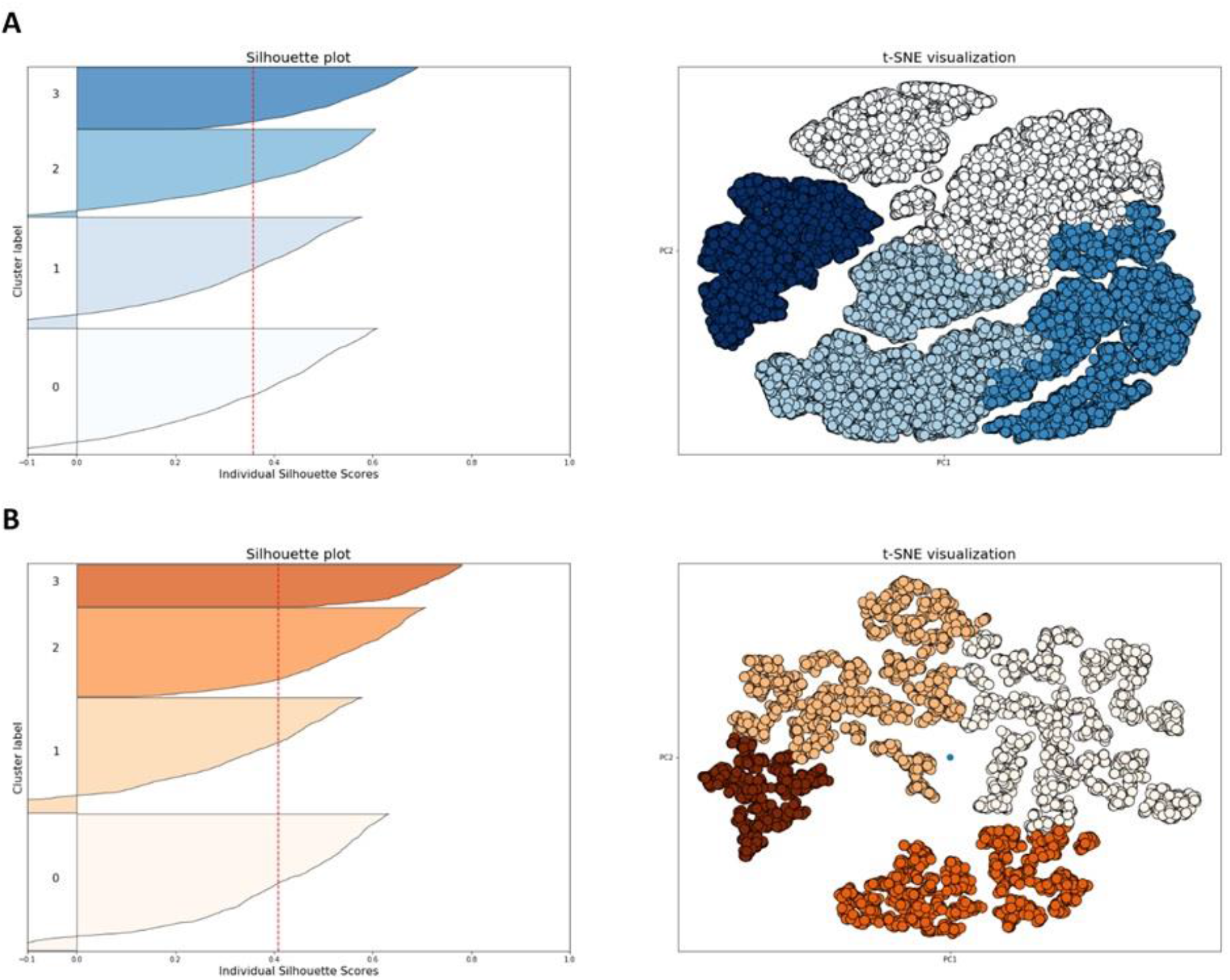
Internal validation and clustering. A) Untreated silhouette score and t-SNE plot. 33 350 datapoints on a t-SNE plot with a perplexity of 75 in four clusters using agglomerative clustering with Ward’s linkage, giving a silhouette score of 0.36. (B) Treated silhouette score and t-SNE plot. 3 979 datapoints on a t-SNE plot with a perplexity of 25 in four clusters using agglomerative clustering with Ward’s linkage, giving a silhouette score of 0.41.

**Table 2.** Cluster summaries and interpretation labels for evaluation. ‘Count’ is the total number of discrete phases in each cluster; ‘Count PCT’ is percentage of phases; ‘Total Duration’ is the total time, in seconds, for each group. ‘Duration PCT’ is percentage of duration. ‘Mean Speed’ and ‘Mean SD’ are group means given in mm/s. ‘Mean Autocorrelation’ and ‘Mean Duration’ are group means. An autocorrelation of 1.0 represents a perfect correlation and 0.0 represents no correlation.

| Label | Untreated |  |  |  | Treated |  |  |  |
| --- | --- | --- | --- | --- | --- | --- | --- | --- |
|  | Swoop | Approach | React | Walk | Swoop | Approach | React | Walk |
| Count | 5 332 | 7 572 | 10 850 | 9 596 | 924 | 1 198 | 1 419 | 438 |
| Count PCT (%) | 15.99 | 22.70 | 32.53 | 28.77 | 23.22 | 30.11 | 35.66 | 11.01 |
| Duration (s) | 24 625 | 46 918 | 63 280 | 56 534 | 33 326 | 8 259 | 8 287 | 3 241 |
| Duration PCT (%) | 12.87 | 24.52 | 33.07 | 29.54 | 14.39 | 35.73 | 35.85 | 14.02 |
| Mean Speed (mm/s) | 301.06 | 92.80 | 122.49 | 51.42 | 369.17 | 90.23 | 182.32 | 33.59 |
| Mean SD (±) | 131.40 | 93.46 | 181.82 | 93.83 | 139.05 | 102.54 | 107.16 | 63.93 |
| Mean Autocorrelation | 0.17 | 0.03 | 0.04 | 0.02 | 0.27 | 0.03 | 0.08 | 0.02 |
| Mean Duration (s) | 4.62 | 6.20 | 5.83 | 5.89 | 3.60 | 6.89 | 5.84 | 7.40 |

### Interpreting results

Mean statistics of each group were investigated to interpret the results (Table 2). Similarly broad behavioural types were found in both arms of the study. After interpretation, these groups were labelled ‘swooping’, ‘approaching’, ‘reacting’ and ‘walking.’ ‘Swooping’ captures fast, short and highly autocorrelated movements; ‘approaching’ slower, less variable behaviour with low autocorrelation; ‘reacting’ faster, more variable actions with some autocorrelation; and ‘walking’ encompasses long, slow movements that are not autocorrelated. As only information about vector movements is used in classification, environmental interactions (e.g., net contact) cannot be included in the definition of behaviours. The labels, and the broad nature of each grouping, were then confirmed through investigation of representative samples from each group.

### Conclusions

Seven principal conclusions can be drawn from this analysis into *An. gambiae* s.s. activity around untreated and treated nets: (1) in any fixed time period, mosquito flight activity is significantly greater when the human host is protected within an untreated net compared to a treated net (two sample Z-test, P < 0.01); (2) four behavioural modes are exhibited around both treated and untreated nets; (3) The proportion of ‘swooping’ behaviour increases significantly around a treated net (two sample Z-test, P < 0.01); (4) The proportion of ‘approaching’ increases significantly in the presence of a treated net (two sample Z-test, P < 0.01); (5) The proportion of ‘reacting’ increases significantly around a treated net (two sample Z-test, P < 0.01); (6) The proportion of ‘walking’ decreases significantly around a treated net (two sample Z-test, P < 0.01); and (7) *An. gambiae* s.s. ‘swoop’ faster in experiments with a treated net (GLM, P < 0.001).

### External Validation

A similar study of the effect of bednet treatment on vector behaviour had previously been conducted in Tanzania using wild *An. arabiensis*, a sibling species closely related to *Anopheles gambiae s.s*. and that exhibits many of the same host seeking behaviour characteristics [9]. This previous study used expert knowledge to identify behaviour types, determining that mosquitos exhibited four behaviours around both an untreated and treated net (‘swooping’, ‘visiting’, ‘bouncing’ and ‘resting’) and that total activity levels dropped significantly at ITNs compared to untreated bednet (from a geometric mean time of 73.5 mins to 23.8 mins). Where particular behaviours are concerned, and comparing total mean times, the study found that ‘swooping’ (where “tracks do not contact the bednet”), ‘visiting’ (“long periods of flight are interspersed with infrequent net contacts’) and ‘resting’ (“mosquito movement is under 1.33 mm /s”) all increased in the presence of a treated net (however, this increase in swooping was not found to be statistically significant). The study also found that ‘bouncing’ (“rapid contacts with the bednet surface… include[ing] walking”) reduced significantly around the treated net (when evaluating geometric mean times).

Comparing findings from this study and those of [9], several replications are clear: (1) both studies recognised four types of mosquito behaviour; (2) total vector activity fell at a treated bednet; (3) ‘swooping’ behaviour increased with a treated net; and (4) ‘walking’ / ‘bouncing’ is decreased when using a treated net. However, although ‘swooping’ and ‘bouncing’ from [9] are acceptable analogues to the behaviours ‘swooping’ and ‘walking’ from this study, it is not possible to align the prior study’s ‘resting’ and ‘visiting’ with the ‘approaching’ or ‘reacting’ categories of this study, due to divergent definitions.

As such, comparison with previous results can only be said to validate four of this study’s conclusions (i.e., (1), (2), (3) and (6) from the Conclusions section). Although there are slight differences between [9] and the current study (i.e., in vectors observed and the definition of behaviour modes), these differences are minor, potentially explicable by the use of a wild population, which is inherently more genetically diverse. With this knowledge, the similarities are such that [9] can be said to support several major findings from this study in the given setting. Consequently, the external validity of the new method was deemed to be proven.

## Discussion

In this study, we present an automated, generalised method for the identification and classification of the behaviour of vectors based on their movement trajectories. This new workflow combines BCPA [19,20] and unsupervised machine learning [31,32] and offers a new solution to current challenges faced by vector biologists and for vector control [1–3]. Although a similar methodology has been proposed for the investigation of marine animal behaviour [36], to our knowledge this is the first use of such an approach within entomology. The method has particular relevance in vector biology, where an automated, repeatable and generalisable means of identifying and defining behaviour that has been validated against vector activity is most pertinent.

Here we supply a preliminary application of the new method, analysing the behaviour of *An. gambiae* s.s., in the presence of both baited untreated bednets and baited ITNs. As the study replicated previous findings, the method is deemed to be an innovative, validated and productive approach that improves and expands the existing toolkit available to vector biologists. Furthermore, the method is repeatable, as any individual with the same dataset will produce the same behavioural tokens and behavioural types; it is generalisable, as it is not limited to a single domain, but can be applied to any vector in any setting; and its foundation in mathematical processes ensures it is immune to observer bias. The accuracy of the new method is confirmed using both internal and external validation [37,38]. The former ensures the correct algorithm and initial parameters are applied, while the latter tests the accuracy of the approach itself. Internal validity is measured in two ways: (1) formal metrics of similarity between datapoints (i.e., silhouette scores) are studied; and (2) t-SNE visualisations of cluster assignment are manually inspected [39–41]. External validation is achieved by replicating known results [42]. The new method is used to compare the behaviour of an insecticide susceptible strain of *An. gambiae* s.s. (Kisumu) around both Long-lasting Insecticidal Nets (ITNs) and untreated nets, producing findings that are corroborated by previous research [9].

The new method offers advantages over alternative, objective approaches that are theoretically automated, repeatable and generalisable. One such method, Hidden Markov Models (HMMs) are probabilistic models that determine the underlying hidden states (e.g., behavioural modes) that cause an observed process (e.g., movement trajectories) [43]. However, for HMMs to apply in this instance, a vector’s behavioural states must be a first-order Markov process. That is, a vector’s behaviour at time *t* must be determined solely by their behavioural state at time *t-1* [43–45]. Nevertheless, it is reasonable to assert that vector behaviour is influenced by internal and external drivers acting over greater periods of time than this and that vector ecology is determined by a wider range of datapoints that cannot be described by a first-order Markov process and an HMM [46]. To capture this more nuanced conception of vector behaviour, a sliding window, such as is applied in BCPA, is needed. Similarly, although several path segmentation methods exist for detecting changes in animal movements other than BCPA [47–50], a form of segmentation that can account for the particular difficulties encountered when tracking vectors must be used in this instance. As the key difficulty here is the frequency of lost frames (caused by the recording system momentarily losing track of the small vector), a method that can handle an irregular dataset is required. As BCPA is a likelihood-based form of path segmentation, which sweeps an analysis window over an entire movement path to identify significant shifts in a parameter value, it provides a robust method for dissecting vector activity into behavioural tokens that can account for irregular temporal measurement intervals and does so without any *a priori* assumptions [19,20].

Although offering several advancements, the method presented here is subject to its own limitations. One such constraint concerns the clarity of the silhouette created by any grouping of behaviours. That is, as behavioural units are nebulous concepts, any silhouette of their classification will be equally unclear and datapoints from different behaviours will not necessarily have a high separateness [51–53]. For example, the distinction between fast walking and slow running is not clear. Consequently, the identification of strong patterns when assessing the clustering of behaviour is unlikely. This is shown in the contiguous silhouettes and the silhouette values produced by movement data (Fig 3). Additionally, it is important to make explicit the assumptions on which this study is based. These assumptions are that vectors are always in some behavioural state, that vectors have more than one potential behavioural mode and that these modes are discrete and expressed over a period of time. Finally, it needs to be clarified that the method presented here is not totally objective. Since the workflow’s Interpretation stage requires experts to attach a label to clusters, a level of subjectivity is still required to implement this analysis. Although this labelling is not theoretically necessary to produce and compare results (as clusters can be described by their characteristics alone), a level of subjectivity is still needed to interpret these results and apply them to everyday discourse concerning behaviour [51–53].

In conclusion, we present and test a new workflow that represents an innovative use of path segmentation and unsupervised machine learning to classify vector behaviour and expands the analytic toolkit available to researchers. This represents a promising development that can improve the evidence base available to vector biologists and open new avenues for the exploitation of vector behaviour to improve intervention performance. Given that global vector control is currently facing a raft of challenges – including environmental and species distribution changes [2], limited resources [3] and an increase in insecticide resistance [54] – novel methodological approaches are more important than ever. Furthermore, it is likely that developments can be made to improve performance and applicability. For example, an analysis of transitions between behaviours could be undertaken, potentially providing additional insights into vector activity and ensuring ecological limits to behavioural transitions have been captured. Finally, the output from this workflow could be used as input to a supervised machine learning algorithm, increasing the efficiency of future analyses.

## Materials and Methods

We present a four-stage workflow in which vector movement trajectories are first collected and pre-processed via BCPA. The most appropriate unsupervised clustering algorithm, and initial parameters, are then identified and applied before the workflow concludes with the interpretation of results, decoding and attaching a behavioural label to each group. The whole workflow is then validated by measuring the accuracy of its results.

### Resources

The workflow presented here is implemented using a combination of R and Python. R is used for pre-processing, utilising the BCPA package built for that language. Python, through a Jupyter notebook, is used at the clustering stage to exploit the `scikit-learn` library. We recommend that the Anaconda platform be used to access RStudio and JupyterLab, as up-to-date installations for Windows, Linux and Mac can all be found in that single distribution. Code, and further details, needed to run the workflow can be accessed through a public GitLab repository: https://gitlab.com/MTFowler/lstm_flightcluster. All analysis found here was performed on a ThinkPad X1 Carbon, using an Intel i7-7500u CPU.

All procedures associated with the collection of mosquito flight data are as described in [9,11,55,56]. Briefly, the ‘Kisumu’ laboratory strain of *An gambiae*, a primary malaria vector across sub-Saharan Africa and susceptible to all insecticides, was used in both the untreated and treated arms of the experiment. All mosquito flight assays were completed in a purpose-built climate-controlled insectary in Liverpool.

### Data acquisition, cleaning and assessment

Vector movement paths were represented by spatial identifiers ordered sequentially via a time variable [18–20,30]. Each event was captured by a unique identifier, an x (longitude or easting) coordinate, a y (latitude or northing) coordinate and a time variable (Fig 1A). This data was collected using an optical imaging and flight-tracking system detailed in [55,56]. This system allowed for multiple vectors to move unconstrained within an enclosed area, a subset of this space being within the field of view of the recording system, creating the recording volume. After collection, movement trajectory data was cleaned and assessed (Table 1). Movements considered noise were removed. This ‘noise’ included short tracks deemed to be isolated fragments from a larger track, or disturbance that has been missed during video cleaning [9–11]. Furthermore, although BCPA accounts for semi-regular sampling [19,20], allowing for some irregularity in the dataset, movement tracks were removed from the analysis if they contained two datapoints at the same time or if they had especially large time gaps (i.e., greater than 10 seconds). Finally, as path segmentation assumes that all time series data displays serial dependence, it was confirmed that the dataset was autocorrelated (i.e., that the velocity of each datapoint is statistically correlated with its recent past) [20]. This was accomplished in R using the ‘Autocorrelation and Cross-Correlation Function Estimation’, `ACF()`.

### Path segmentation

With a correctly formatted dataset, that had been cleaned and assessed, BCPA was applied. BCPA is a form of path segmentation that identifies changes in animal behaviour, at the path-level, based on significant shifts in a parameter value of an organism’s movement trajectory. As BCPA accepts movement paths as sequentially ordered step lengths, turning angles and velocities, rather than the spatial identifiers collected by tracking technology, spatial values were converted into the required variables using the `GetVT()` function from R’s `BCPA` package [57]. Within BCPA there are four user defined parameters: (1) the ‘Parameter Value’ (the response time-series variable in which significant changes will identify a behavioural change point); (2) the ‘Window Size’ (the number of datapoints the window will capture when sweeping); (3) a sensitivity parameter ‘K’; and (4) the ‘Cluster Width.’ For arthropod activity, it was determined that optimal segmentation occurs at a significant change in persistence velocity (Velocity*cos(Turning Angle)), using a window size of 30, a window step of 1, a sensitivity value of 2 and a cluster width of 1. These initial parameters were determined following BCPA documentation recommendations [19,20,57] and to maximise sensitivity to behavioural shifts. (Note, however, that this increase in sensitivity amplifies the chances of spurious shifts being detected which will ultimately result in transitions to the same behaviour in the final output. However, as the alternative is to lower sensitivity and potentially miss legitimate changes in behaviour, a high sensitivity, with corollary spurious shifts, is preferred.)

### Clustering

To determine the optimal unsupervised learning algorithm and initial parameters for clustering, internal validation was undertaken. Following [37,38], the form of internal validation used was silhouette scores [58,59] and visual inspection of t-SNE plots [39–41]. A silhouette score measures how well data had been grouped, comparing each object’s similarity to others within its own cluster (group tightness) and those from other clusters (group separation) and was calculated using Python’s `silhouette_score()` function from the `metrics` module of the `scikit-learn` library. This measure gives a score between −1.00 and +1.00, with a silhouette value below 0.20 showing no structure is present in the data and the grouping is invalid; a figure over 0.70 representing a strong structure and a valid grouping; and a silhouette score around 0.50 illustrating that a reasonable structure has been found within the data and that the clustering is acceptable [59]. Detailed silhouette coefficients for each sample was then visualised using a silhouette plot in Python with the `silhouette_samples()` function from the `sklearn.metrics` module (Fig 1C). As all clusterings require manual review to validate appropriateness [37], the high-dimensional data was reduced and positioned in a two-dimension map using t-distributed Stochastic Neighbour Embedding (t-SNE) [39]. Once mapped, the appropriateness of the clustering was verified through manual visual inspection. Review ensured there were acceptable levels of cohesion between members of the same group and separateness between members of different groups. t-SNE was undertaken using the `TSNE()` function from the `sklearn.manifold` module found within Python. When performing t-SNE, the user needs to define the perplexity (an estimate number of nearest neighbours for each datapoint), with larger datasets requiring a larger perplexity [41]. Consequently, perplexity was fine-tuned to show global geometry.

Once the optimum algorithm and parameters were determined by silhouette score and inspection of t-SNE plot, findings were applied to the BCPA output. Python’s `scikit-learn` library was used as it is an efficient means of building standard machine learning models [60]. (Other packages, such as R’s `class`, are available when implementing unsupervised machine learning, however different packages may produce different results.)

### Interpreting results

The final stage of analysis is to interpret and label clusters [35]. Although the naming of individual clusters can be systematised, its final interpretation is ultimately a somewhat subjective process. Interpretation here entailed scrutiny of the attributes of each cluster. Taking each group’s mean velocity, standard deviation, autocorrelation and duration, an expert analysed and named the behaviour associated with the movement classes. This initial understanding was then confirmed and refined through visual inspection of representative samples. ‘Representative’ was defined as those samples found at the centre point of a group and such datapoints were deemed to be the most typical of that behaviour class [33,34]. Consequently, interpretation of results was bolstered through centroid analysis, with those datapoints at the heart of each group’s t-SNE mapping, or as close to the centre as possible, isolated and visually inspected by an expert. Multiple such examples close to the centre were isolated and inspected, thereby confirming, or refining, the initial analysis.

Using both the analysis of descriptive statistics and centroid analysis, an expert was able to interpret the broad behavioural type of each cluster and attach a sensible label. If no label was able to be attached, either because no behaviour is being demonstrated or behaviours are spread between clusters, the full workflow was undertaken again. By manipulating the user defined settings during BCPA or dimensionality reduction, significant changes in clustering can result. Consequently, these parameters were fine-tuned to optimise performance and final behaviour identification.

### External Validation

To ascertain the accuracy of the new method, its performance was externally validated by comparing results concerning the difference in flight patterns of *An. gambiae* s.s. around human-baited insecticide-treated bednets (ITNs) and untreated bednets. Findings generated by the new, computation-derived, method were contrasted with those from a previous study that employed the existing, expert-defined, method for behavioural classification. Although a standard method of external validation is through comparison against an *a priori* dataset, testing whether a known, true, clustering can be recreated [38], this was not possible in this instance. There is no known, incontrovertibly true, grouping for this *An. gambiae* s.s. behaviour and therefore no such comparison can be made. Consequently, external validation was established via confirmation with previous results. That is, the workflow’s accuracy as a method was corroborated by comparing its conclusions to those already present in the literature [9] to verify whether prior findings could be replicated.

## Acknowledgements

We are grateful to Elizabeth Bandason, Amy Guy and Josie Parker for advice on interpreting mosquito behaviour and James Maas for advice on statistical methods.

## Author Contributions

Conceptualisation, formal analysis, methodology and writing (draft preparation) by M.F. Data curation and writing (review & editing) by A.A. and G.M. Software by V.V., C.T. and D.T. Supervision and writing (review & editing) by P.M. All authors have read and agreed to the published version of the manuscript.

## Data availability statement

All data and code used for running experiments, model fitting, and plotting is available on a GitLab repository at https://gitlab.com/MTFowler/lstm_flightcluster.

